# Interleukin-17A Secreted from the Lung–infiltrating T Helper 17 Cells Renders Protective Immunity to Pulmonary *Cryptococcus neoformans* Infection

**DOI:** 10.1101/328005

**Authors:** Elaheh Movahed, Grace Min Yi Tan, Heng Choon Cheong, Chalystha Yie Qin Lee, Yi Ying Cheok, Sun Tee Tay, Pei Pei Chong, Won Fen Wong, Chung Yeng Looi

## Abstract

IL-17A has emerged as a key player in the pathologies of inflammation, autoimmune disease, and immunity to microbes since its discovery two decades ago. In this study, we aim to elucidate the activity of IL-17A in the protection against *Cryptococcus neoformans*, an opportunistic fungus that causes fatal meningoencephalitis among AIDS patients. For this purpose, we examined if *C. neoformans* infection triggers IL-17A secretion in the *in vitro* setting using RAW264.7 murine macrophage cells, and *in vivo* using wildtype C57BL/6 mice. In addition, an enhanced green fluorescence protein (eGFP) reporter and a knockout (KO) mouse models were used to track the source of IL-17A secretion and explore the protective function of IL-17A, respectively. Our findings showed that both *in vivo* and *in vitro* models of *C. neoformans* infection demonstrated induction of abundant IL-17A secretion. By examining the lung bronchoalveolar lavage fluid (BALF), mediastinal lymph node (mLN) and spleen of the IL-17A– EGFP reporter mice, we showed that intranasal inoculation with *C. neoformans* promoted leukocytes lung infiltration. A large proportion (~50%) of the infiltrated CD4^+^ helper T cell population secreted EGFP, indicating vigorous T_H_17 activity in the *C. neoformans*–infected lung. The infection study in IL-17A–KO mice, on the other hand, revealed that absence of IL-17A marginally boosted fungal burden in the lung and accelerated the mouse death. Therefore, our data suggest that IL-17A, released predominantly from T_H_17 cells *in vivo*, is essential in providing a protective immunity against *C. neoformans* infection.

## INTRODUCTION

The opportunistic pathogenic basidiomycete *Cryptococcus neoformans* is an encapsulated yeast commonly found in bird excrement worldwide (1). The infection is often asymptomatic in healthy individuals but causes severe pulmonary cryptococcosis and life-threatening meningoencephalitis in immunocompromised patients. *C. neoformans* has gained attention in recent years as it is a major cause of death among patients who have advanced acquired immunodeficiency syndrome (AIDS) (2). Hence, it is important to study the host interaction with this pathogen as 30-60% of the patients who have cryptococcal meningitis succumb to cryptococcosis infection within one year despite antifungal therapy (3).

The early interaction between innate immune cells and cryptococci is critical in determining the outcome of cryptococcosis (4). The presence of *C. neoformans* cells or its major capsular polysaccharide glucuronoxylomannan promotes NF-κB nuclear translocation and activation in the macrophages (5), which subsequently induces secretion of cytokines including tumor necrosis factor alpha (TNF-α) and transforming growth factor β (TGF-β) (6). Activated macrophage or other phagocytes are able to eliminate most of the engulfed pathogens via formation of phagolysosome, nitric oxide and reactive oxygen activities. As with other successful pathogens such as *Helicobacter pylori, C. neoformans* has evolved a number of elaborate strategies to evade immune destruction by macrophages *(7)*. The encapsulated *C. neoformans* can defend itself against the onslaught of macrophages through non-lytic expulsion (8, 9), a process that involves alteration of host cell Arp2/3 complex-mediated actin polymerization and phagosome pH (10, 11), thereby allowing the pathogen to survive and propagate within macrophages. *C. neoformans-*activated macrophage often displays an M1-like phenotype (12) whose polarization is highly plastic depending on external signals such as cytokines (13). *C. neoformans* intervenes in the polarization of macrophages by using Ssa1, a heat shock protein 70 homolog to drive macrophage development toward the alternative M2 phenotype that is defective in the fungal clearance (14). These processes impair the host innate immune system and promote the invasion and dissemination of *C. neoformans* in the host.

The essential role of T cells in the host immune response to *C. neoformans* has been well-studied using T cell depletion mouse model (15-17). In the era of two distinct helper T (T_H_) paradigm, T_H_1 or T_H_2, most findings are in agreement that protective immune response against the fungus is principally driven by T_H_1 cells, whereby IL-12 and IL-18 potentiate T_H_1 polarization and interferon-γ (IFN-γ) and TNF-α release which contribute to fungal clearance (18-20). Some studies suggest the involvement of T_H_2 as IL-4 is detected in high quantity in mice infected with highly virulent strain at 2 to 3 weeks post infection (21, 22). A dominant T_H_2 cytokine profile has been associated with increased cryptococcal proliferative potential (23). IL-4 and IL-13 released by T_H_2 or eosinophils in lung could also cause fatal allergic inflammation during bronchopulmonary mycosis (24, 25), a reaction which is dampened by IL-23 through an IL-17A–independent and –dependent mechanisms (26). Following the recent breakthrough in T cell subsets discovery, a relatively new T cell subset, T_H_17, has also been implicated in the immune response to fungus (27, 28). A study has also challenged the role played by IL-17A in anti-fungal response and claimed that IL-17A promotes the fungal infection (29), as such, the nature and role of these T cell subsets require further investigations. T_H_17 cell is characterized by its hallmark RORγt transcription factor and IL-17 secretion. Its differentiation from naïve CD4^+^ T cells is induced in the presence of IL-6 and TGF-β during inflammatory response. IL-23 is another important inducer for IL-17A as the IL-17A production was strongly impaired in the IL-23p19 deficient mice (30). *C. gattii*, a highly virulent cryptococcal species is able to attenuate both T_H_1 and T_H_17 by suppressing *IL-12* and *IL-23* genes transcription (31).

T_H_17 is not the sole source of IL-17A as it can also be released by other cells such as macrophages, NK cells, and neutrophils (32). IL-17A elicits inflammatory response by recruiting neutrophils, but does not contribute to classical macrophage activation as seen in pulmonary cryptococcosis induction in the mouse model (33). T_H_17 cells release a panel of other cytokines in addition to IL-17 such as IL-17F, IL-22, and IL-23. A full picture of regulatory mechanism as to how this subset of T cell interacts and eliminates the fungal infection requires further investigation. In this study, we examined the association of IL-17A with *C. neoformans* infection by using both *in vitro* and *in vivo* infection models. The main focus of our study lies on identifying the source of IL-17A secretion and determining its protective role in *C. neoformans*– infected mice. Using enhanced green fluorescence protein (eGFP) reporter mouse model, we showed that lung infiltrating T_H_17 cells are likely the predominant source of IL-17A. Data from a knockout (KO) mouse model supports a protective function of the IL-17A against *C. neoformans* infection.

## RESULTS

### *C. neoformans* infection induces IL-17A production in both *in vitro* and *in vivo* models

To examine if *IL-17A* gene transactivation is induced by *C. neoformans* infection, RAW264.7 mouse macrophage cells were first cultured with *C. neoformans* H99 strain at MOI 5:1 for 24 hours followed by quantitative real-time PCR. Intriguingly, the relative expression level of IL-17 to β-actin was drastically increased at 4.4–fold in the *C. neoformans*–infected cells compared to the non–infected control (Fig. 1). The relative expression levels for other cytokines, i.e. IL-6 and IFN-γ were also increased at 3.2– and 2.0–fold respectively, indicating initiation of pro-inflammatory response along with T_H_1– and T_H_17–inducing cytokines upon *C. neoformans* infection.

**FIG 1.**
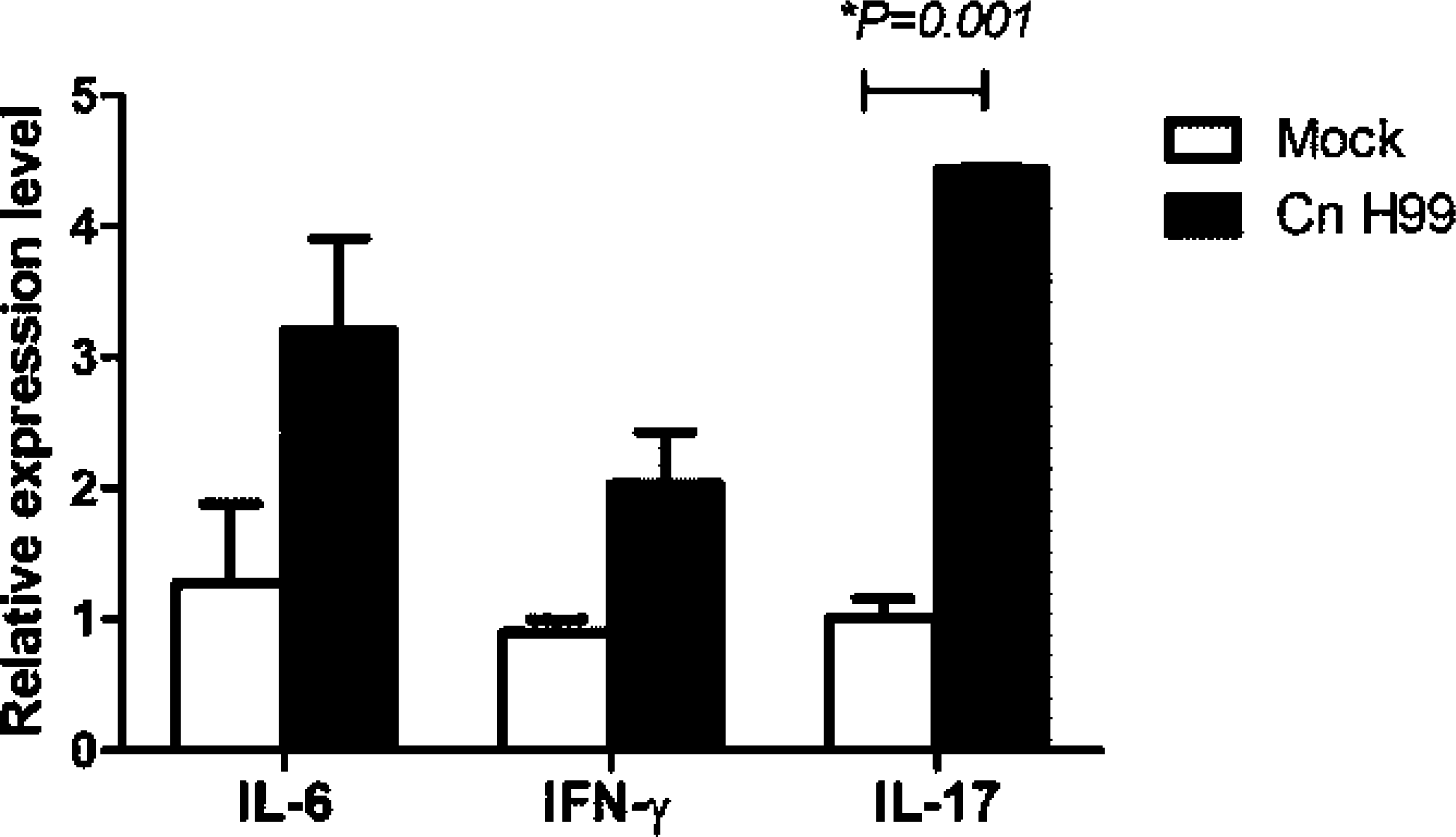
Increased expression of cytokines in *C. neoformans*–infected macrophages. Quantitative real-time PCR result shows Relative expression levels of IL-6, IFN-γ and IL-17 mRNA transcripts of the RAW264.7 cells uninfected (mock) or infected with *C. neoformans H99* (Cn H99) for 24 hours. Data is mean ± SD from one experiment run in triplicate, and is representative of two independent experiments. \**P*<0.05 by Student’s *t*-test.

To investigate if *C. neoformans* infection–mediated IL-17A secretion occurs *in vivo*, C57BL/6 mice were intranasally inoculated with four different strains of *C. neoformans* (H99, S48B, S68B and H4) at 2 × 10^5^ cells for 14 days before collecting serum for cytokine measurement (Fig. 2). Consistent with the data from *in vitro* infection, an elevated serum level of IL-17A was detected in the *C. neoformans*–infected mice (Fig. 2A). We noted that the serum IL-17A level was correlated with the degree of virulence of different *C. neoformans* strains (34), whereby the highest amount of serum IL-17A was observed in the group of mice infected with the most virulent *C. neoformans* H99 strain (115 ± 12 pg/ml). This was followed by moderate serum IL-17A level observed in the mice infected with less virulent environmental strains, S48S (89 ± 3 pg/ml) and S68B (75 ± 2 pg/ml), and lowest level of serum IL-17A was noted in the mice infected with non-virulent strain H4 (24 ± 1 pg/ml) strains, compared to control uninfected mice (<20 pg/ml). The level of serum IL-23 was also elevated in all *C. neoformans*–infected mice, i.e. H99 (67 ± 5 pg/ml), S48S (78 ± 3 pg/ml), S68B (36 ± 1 pg/ml) and H4 (47 ± 5 pg/ml) compared to <20 pg/ml in the mock control (Fig. 2B). This suggests IL-23–IL-17A axis pathway plays a major role in the host immunity against *C. neoformans* infection. On the other hand, the serum IL-17F levels were only scarcely increased in mice infected with *C. neoformans* H99 and S48B strains (Fig. 2C), while no noteworthy induction was observed for other cytokines (MIP-3α, IL-21, IL-31 and IL-33) examined (data not shown).

**FIG 2.**
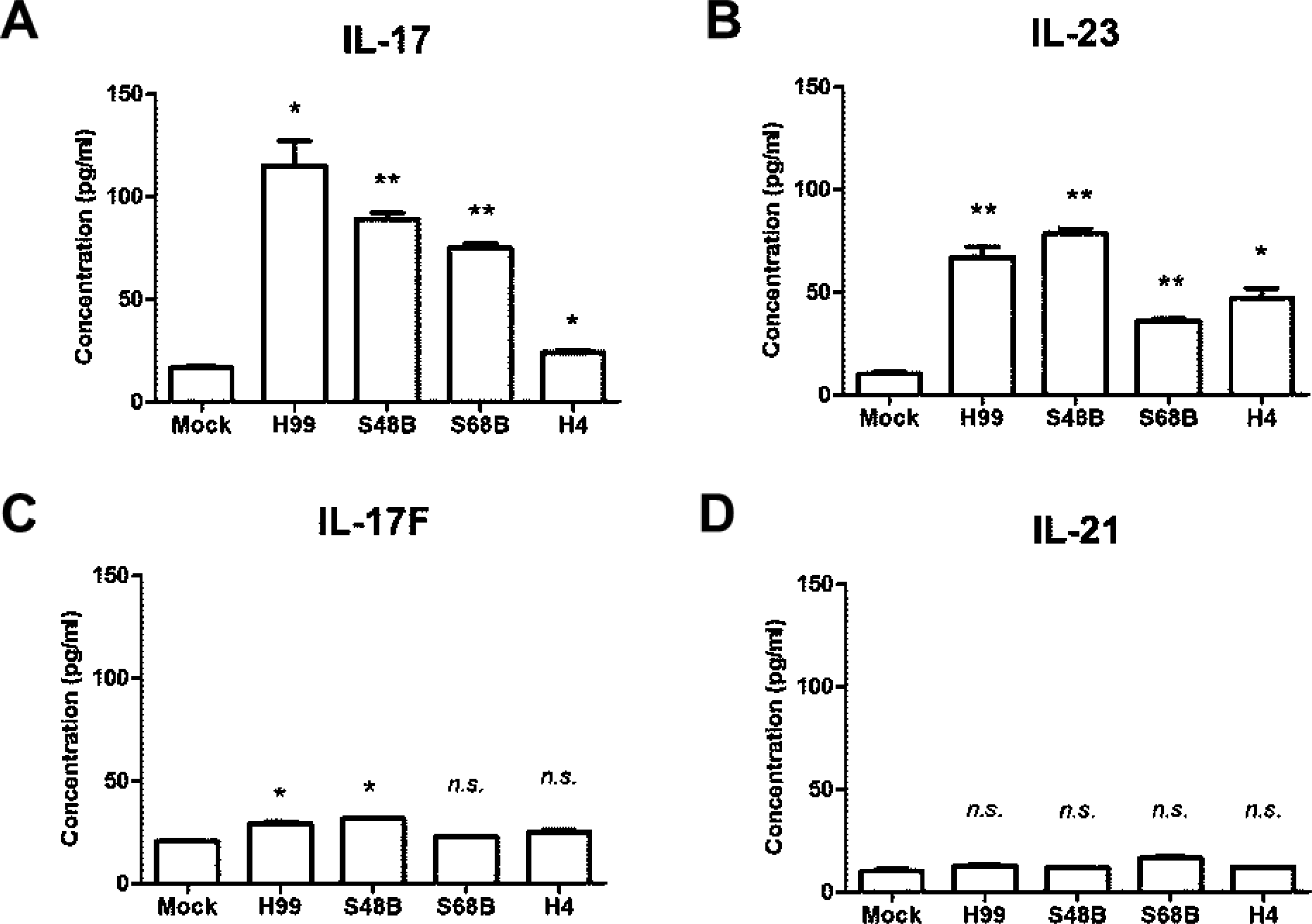
Elevated serum IL-17A, IL-23 and IL-17F levels in the *C. neoformans*–infected mice. C57BL/6 mice (n=4 per group) were intranasally inoculated with 1 × 10^5^ cells of four different strains of *C. neoformans* (H99, S48B, S68B and H4), serum were collected after 14 days for Bioplex cytokine array. Mock denotes control mice intranasally administrated with equal volume of PBS. Different cytokines in the T_H_17 panel, IL-17A (A), IL-23 (B), IL-17F (C), and IL-22 (D) were measured. \**P*<0.05, \*\**P*<0.01, *n.s.*: not significant or *P*≥0.05, by Student’s *t*-test.

### Intranasal *C. neoformans* inoculation causes leukocytes lung infiltration

To examine the importance of IL-17 in providing immunity to *C. neoformans* infection, we utilized a mouse model harboring IRES-EGFP-SV40-polyA signal sequence cassette after the stop codon of *Il-17a* gene in which the EGFP is co-expressed in the IL-17A–producing cells. Mice were intranasally administrated with 20 μl of control PBS or *C. neoformans* (2 × 10^5^ cells) in suspension, and splenic, mediastinal lymph nodes (mLN), bronchoalveolar lavage fluid (BALF) cells were inspected after 4 weeks (Fig. 3). Total numbers of cells were significantly increased in the BALF (4.8 × 10^6^ ± 1.0 × 10^6^ versus 4.6 × 10^4^ ± 1.0 × 10^4^ cells, \**P*=0.0135) and mLN (3.0 ×10^5^ ± 2.0 × 10^4^ versus 1.5 × 10^5^ ± 1.9 × 10^4^ cells, \*\**P*=0.0017) of the *C. neoformans* H99– infected mice versus the control group (Fig. 3A and 3B). No significant increased numbers of cells were observed in the spleen after intranasal *C. neoformans* H99 infection (5.6 × 10^7^ ± 7.1 × 10^6^ versus 4.7 × 10^7^ ± 6.1 × 10^6^, *P*=0.46), suggesting localized infection (Fig. 3C).

**FIG 3.**
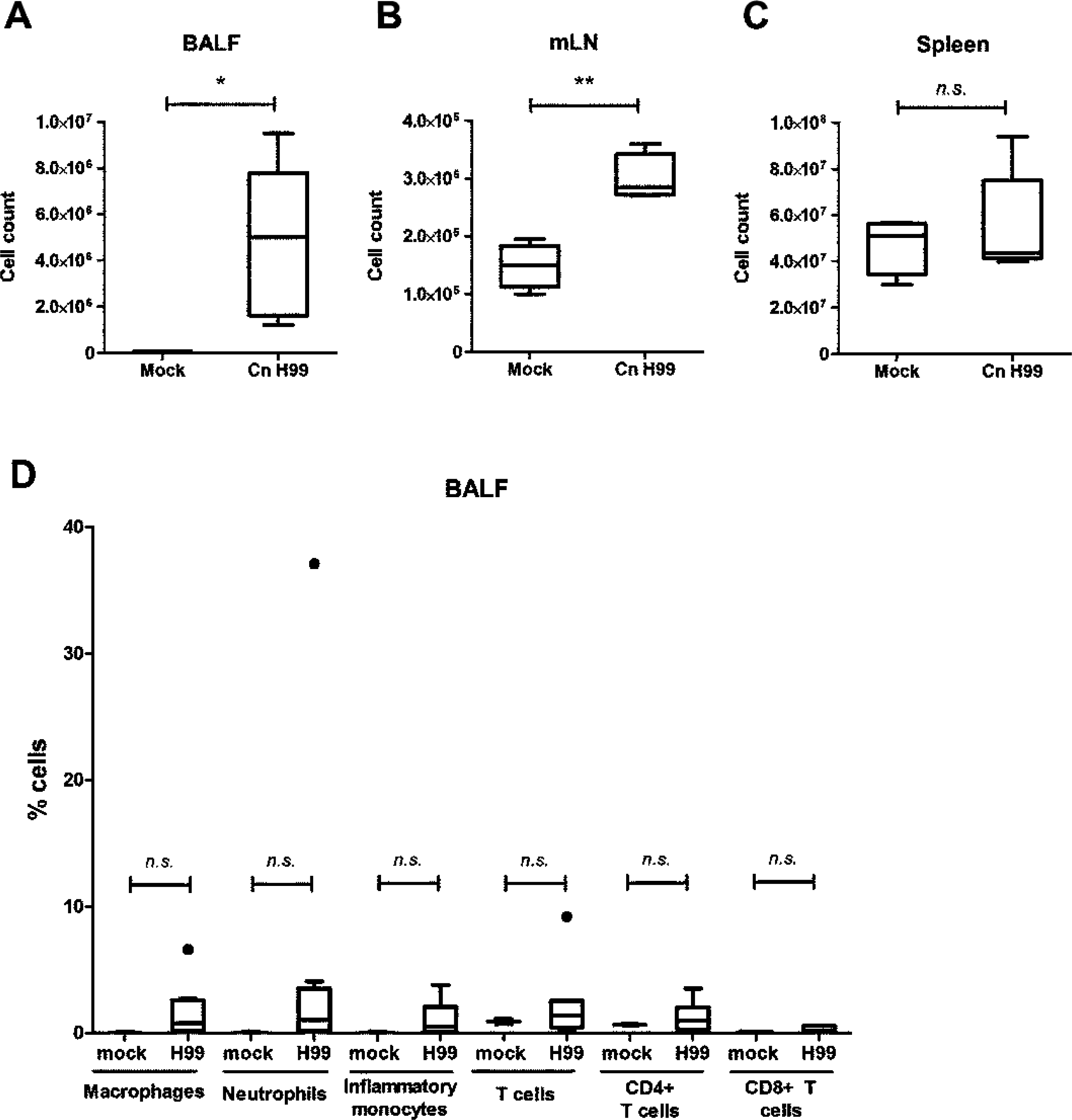
Increased number of leukocytes in BALF and mLN of the *C. neoformans*–infected mice. IL-17A–EGFP reporter mice (n=4 per group) were uninfected (mock) or intranasally inoculated with 1 × 10^5^ cells with *C. neoformans* H99 strain (Cn H99), BALF, mLN and spleen were collected after 14 days for analysis. (A–C) Total number of cells in BALF, mLN and spleen. (D) Total numbers of different leukocytes after examination by flow cytometry analysis. \**P*<0.05, \*\**P*<0.01, *n.s.*: not significant or *P*≥0.05, by Student’s *t*-test.

In the BALF from the *C. neoformans*–infected mice, active immune response was noted as all types of leukocytes examined, except CD8^+^ T cells demonstrated an average of 2-fold increment (Fig. 3D). No significant differences of the cell constituents were observed in lymph node (data not shown). In spleen, the percentages of innate immune cells i.e. macrophages (CD11b^+^F4/80^+^), neutrophils (CD11b^+^Gr1^hi^) and inflammatory monocytes (CD11b^+^Gr1^int^) were increased at approximately 3– to 6–fold. On the contrary, percentages of T (both CD4^+^ and CD8^+^) cells were slightly reduced at 1.2– to 1.3–fold (data not shown).

### Increased IL-17A–producing T cells in the lung of *C. neoformans H99*-infected mice

In the GFP in IL-17A–EGFP reporter mice, IL-17A–producing cells can easily be identified as they display EGFP fluorescence, hence this mouse model was utilized to determine the main source of IL-17 during *C. neoformans* infection. Our result showed that there was no profound increase of GFP^+^ cells amongst macrophage, neutrophil or inflammatory monocytes populations in the *C. neoformans*–infected mice (data not shown). On the contrary, significant increases of GFP^+^ cells were observed among the T cells (Fig. 4). Major source of IL-17A was derived from CD3^+^CD4^+^ but not CD3^+^CD8^+^ T cells. Almost half (54.2 ± 11.6 %, \*\*\**P*=0.0008) of the total lung infiltrated CD3^+^CD4^+^ T cells recovered in BALF collected from *C. neoformans* H99-infected mice were GFP^+^, compared to only 4.8 ± 0.8% in the control (Fig. 4A). In the mLN, the percentages of GFP^+^ CD3^+^CD4^+^ cells were approximately 4–fold greater at 21.7 ± 1.4 % (*P*<0.0001) cells, compared to 5.1 ± 0.6 % in control mice (Fig. 4B). The percentages of GFP^+^ cells among CD3^+^CD4^+^ population were also marginally increased (17.53 ± 4.1 %, *P*=0.016) among the splenic CD4^+^ T cells compared to 6.1 ± 0.6 % in the control (Fig. 4C).

**FIG 4.**
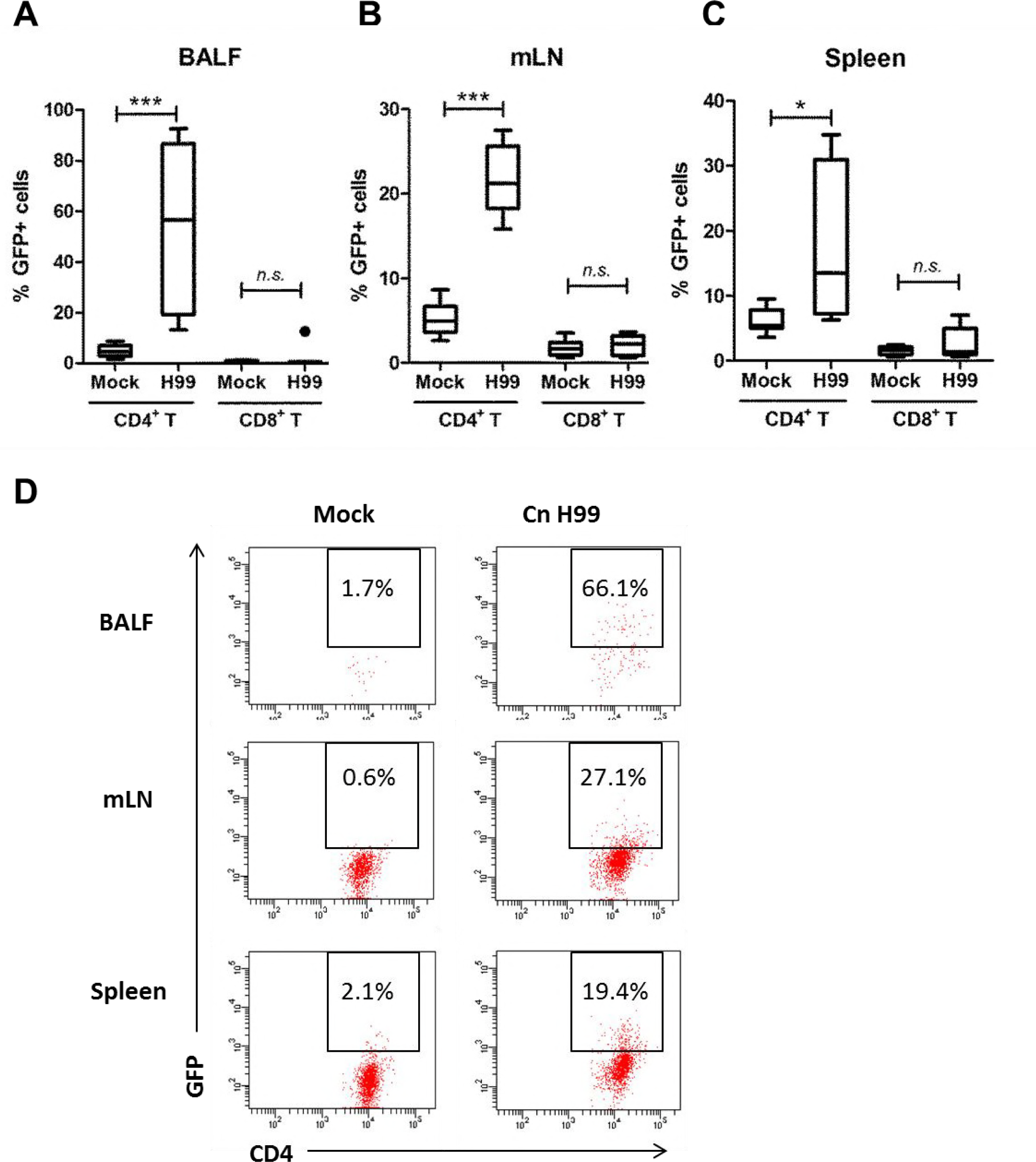
Production of IL-17A by CD4^+^ T helper cells. IL-17A–EGFP reporter mice (n=4 per group) were uninfected (mock) or intranasally inoculated with 1 × 10^5^ cells with *C. neoformans* H99 strain (Cn H99), BALF, mLN and spleen were collected after 14 days for analysis. (A–C) Number of GFP^+^ cells among the CD4^+^– or CD8^+^–gated T cell populations in the BALF, mLN and spleen. \**P*<0.05, \*\**P*<0.01, \*\*\**P*<0.001, *n.s.*: not significant or *P*≥0.05, by Student’s *t*-test. (D) A representative flow cytometrical plot of GFP^+^ cells in BALF, mLN and spleen among CD4^+^–gated T cells. % denotes the percentage of GFP^+^ cells appear inside the gated area.

### IL-17 provides protective immunity to pulmonary *C. neoformans* infection

A knock out mouse model was then applied to examine the protective role of IL-17 to host during *C. neoformans* infection. Intranasal inoculation of *C. neformans* in wildtype C57/BL6 mice and IL-17A–KO resulted in mice death starting from day 26 and 24, respectively. In IL-17A-KO mice, more than 80% (5 out of 6) mice died on day 34 whereas in wildtype control, this was observed at day 40 (Fig. 5A). The amounts of fungal cells in the local infection site (lung) as well as systemic infection (brain) were assessed. CFU counts in the lung derived from IL-17KO mice stayed at 657 ± 92, a higher level compared to 473 ± 119 in the wildtype control, whereas CFU count in the brain was 237 ± 39 compared to 133 ± 30 in the control mice (Fig. 5B). Hence, the absence of IL-17A accelerated mice death as a result of an increased CFU count, suggesting its protective role against *C. neoformans* invasion.

**FIG 5.**
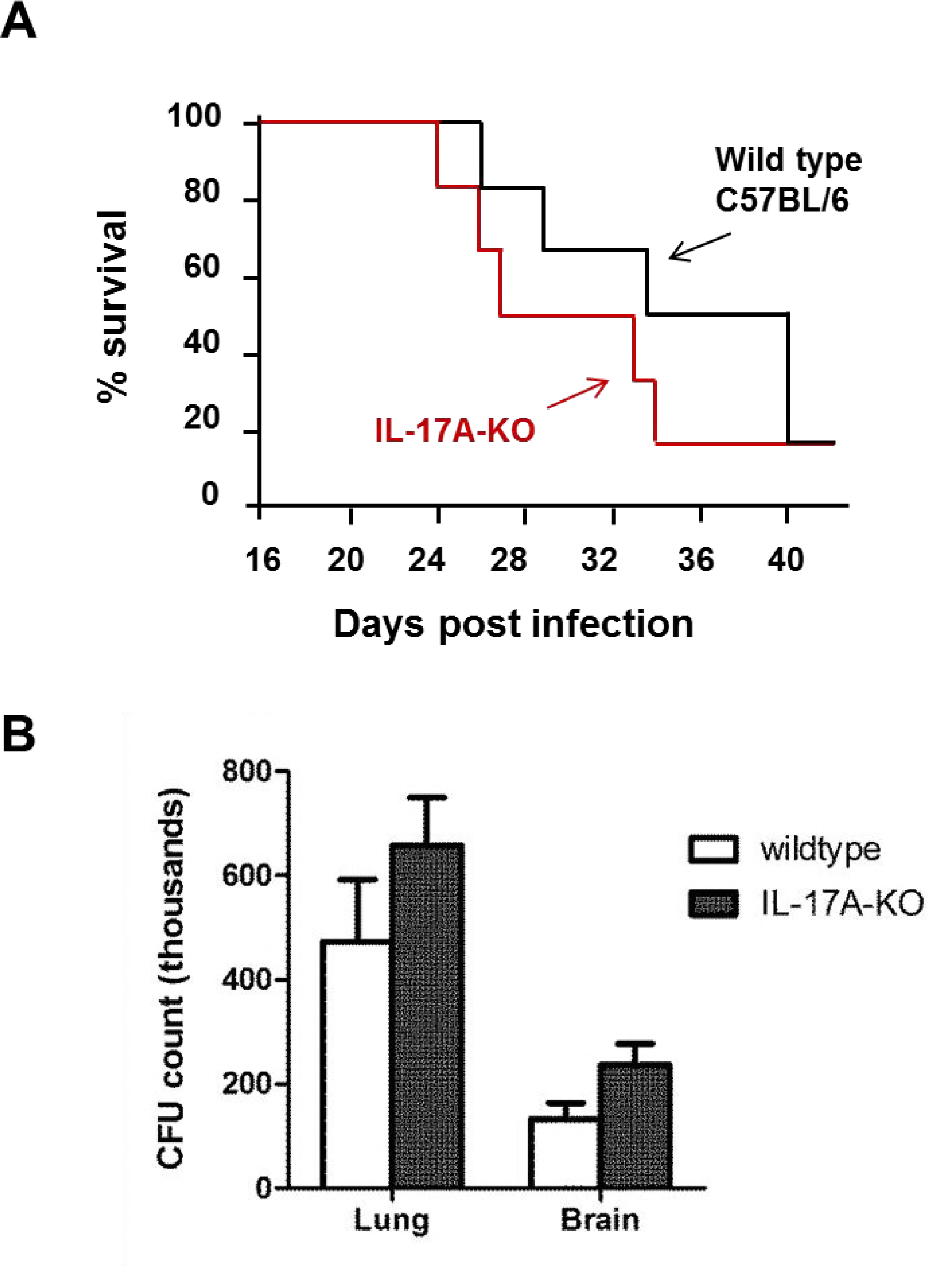
Attenuated protective immunity to *C. neoformans* in the IL-17A deficient mice. (A) Survival curve of the control and *C. neoformans*–infected mice. IL-17A–KO mice (n=6 per group) were uninfected (mock) or intranasally inoculated with 1 × 10^5^ cells with *C. neoformans* H99 strain (Cn H99) and were observed closely over a peroid of 40 days. (B) Fungal burden of the control and *C. neoformans*–infected mice. IL-17A–KO mice (n=5 per group) were uninfected (mock) or intranasally inoculated with 1 × 10^5^ cells with *C. neoformans* H99 strain (Cn H99). Fungal CFU counts in the lung were quantitated after 14 days. Data is shown as mean ± SD.

## DISCUSSIONS

Increasing prevalence of mortality attributed to cryptococcal meningitis in the immunocompromised patients underscores the importance of elucidating the host defense– pathogen interaction. In this study, we focused on (i) determining the expression and source of elevated IL-17A from different types of immune cells using a IL-17A–GFP reporter animal model and, (ii) investigating the role of IL-17A in protective immunity using a IL-17A–KO mouse model. Our data demonstrated elevated serum levels of T_H_17 cytokines, i.e. IL-17A and IL-23 in the infected wild-type C57BL/6 mice; and proposed the lung–infiltrating T_H_17 subset as the major source for IL-17A secretion at the lung upon pulmonary *C. neoformans* infection. In addition, we also showed that the absence of IL-17A resulted in a greater fungal burden and accelerated death of the infected mice, which implies a protective role of the potent IL-17A response at an early stage of *C. neoformans* infection.

Although a high level of IL-17A was detected in the *in vitro* infection model using RAW264.7 macrophage cell line in this study, similar level of expression was not observed in the macrophages of *in vivo* animal infection model with the eGFP reporter system. Classical macrophage activation is not affected in the IL-17A–depleted mice upon pulmonary cryptococcal stimulation, hence IL-17A may not be derived from the activated macrophages (33). Instead, other cells like neutrophils and T cells may serve as the predominant leukocytic source of IL-17A (35). Human monocytes derived from healthy donors exhibited an extensive modification of transcriptome level upon incubation with *C. neoformans*, in particular the genes related to TNF-α, NF-κB, Jak-Stat and toll-like receptors pathways (36). Pro-inflammatory cytokine IL-1β is one of the top up-regulated cytokines in the *C. neoformans*–infected macrophages, and a deficiency in IL-1R1 results in defective T_H_17 cytokines secretion (37). This suggests that in the *in vivo* condition, macrophages may not directly release IL-17A but could instead be involved in indirect induction of IL-17A secretion from other immune cells through the release of pro-inflammatory cytokines such as IL-1β.

Most studies thus far pinpoint the association of T_H_1-type cytokine responses with protective immunity against pulmonary cryptococcosis (38-40). The cytokines in response were mainly those of T_H_1 subsets (IFN-γ, TNFα, IL-8), whereas moderate increases were also observed in T_H_2 (IL-4) and T_H_17 cytokines (IL-17A) (41). A predominant T_H_1 and/or T_H_17 cytokine profile limits the growth of *C. neoformans* and *C. gattii*, whereas a T_H_2 cytokine profile promotes intracellular fungus proliferation (23). In humans, it has also been reported that cryptococcal-specific CD4^+^ T-cell response is predominantly a T_H_1 type response with minimal involvement of T_H_2 and T_H_17 cells (41). However, patients with higher IFN-γ or TNF-α production showed greater level of IL-17A level in their cerebrospinal fluid (CSF). These patients demonstrated lower fungal burdens and faster clearance of *C. neoformans* infection, suggesting that both T_H_1 and T_H_17 responses worked cooperatively to provide optimal immunity against pulmonary cryptococcosis.

Our findings from IL-17A-KO mice experiment are in concordance with previous studies which showed that pulmonary fungal burden was resolved at a slower rate, and the overall survival was not deteriorated due to IL-17A deficiency (33, 35). Besides, higher fungal dissemination to the brain was also observed in the surviving IL-17A–depleted mice, consistent with a previous finding (35). These data support that IL-17A participates in providing protective anti-cryptococcal host defenses through the suppression of fungal growth and dispersal. This observation is in line with other studies on several other fungal species (42-44). It was shown that a deficiency in IL-17A response results in increased susceptibility to oropharyngeal and disseminated candidiasis (27, 45). Decreased neutrophil infiltration, increased fungal burden, and exacerbated pathology were reported upon IL-17A neutralization in *C. albicans* and *Aspergillus fumigatus* infections (42-44). Toll IL-1R8 (TIR8), another negative regulator of T_H_17 response, has also been shown to reduce the susceptibility and immunopathology to candidiasis (46). Some studies, on the contrary, provide evidence that outcome of aspergillosis in human is independent of T_H_17 responses (29), and the IL-23/IL-17A–driven inflammation could impede antifungal immune resistance and promote infection of *A. fumigatus* (47). Hence, further investigation is necessary to validate the precise function of T_H_17 immunity towards fungal infection in humans.

In summary, our data suggest that IL-17A derived from the lung infiltrating T_H_17 in BALF and mLN, plays a supportive role in rendering protection to pulmonary *C. neoformans* infection. Understanding the host immune response during cryptococcal infection is essential for the development of immunomodulatory therapies.

## MATERIALS AND METHODS

### Fungal and cell culture

*C. neoformans var. grubii* (serotype A) H99 and RAW264.7 murine monocytic macrophage cell-line were obtained from American Type Culture Collection (ATCC). Environmental strains S48B, S68B and H4 were isolated from pigeon droppings, as described (48, 49). To start the culture, a small drop of fungal cell stock was streaked on the Sabouraud’s dextrose agar (SDA) and incubated at 37°C for 48 hours. Then, 2 to 3 single colonies from freshly prepared agar plate were inoculated into Sabouraud’s dextrose broth (SDB) and incubated at 37°C for 48 hours. RAW264.7 cells were cultured in Dulbecco’s Modified Essential Medium (DMEM) supplemented with 10 % fetal bovine serum and incubated at 37°C, 5% CO_2_. For *in vitro* infection, RAW264.7 cells were seeded at 5.0 × 10^5^ cells/ml and infected with *C. neoformans* at multiplicity of infection (MOI) of 5 for 24 hours.

### Quantitative real-time PCR

RNA was isolated from cells as described previously (7). Briefly, 1 ml TRIzol Reagent (Invitrogen, Carlsbad, CA) and 200 μl chloroform were added to cells and vortexed vigorously for 15 sec. The upper phase was collected and precipitated with isopropanol, washed with 70% ethanol and dissolved in RNase-free water. Then, cDNA was generated using MMLV reverse transcriptase (Life Technologies). Samples were amplified with SsoAdvanced™ SYBR^®^ Green Supermix (Biorad) using StepOnePlus™ Real-Time PCR Systems (Life Technologies) using the following PCR cycling parameters: 95°C for 10 min, 40 cycles at 95°C for 15 sec each, and 60°C for 1 min. Data were analyzed using StepOne software version 2.3. Primer sequences used were *IL-17A* (5’-TCTCCACCGCAATGAAGACC-3’ and 5’-CACACCCACCAGCATCTTCT-3’), *IL-6* (5’-CCTCTGGTCTTCTGGAGTACC-3’ and 5’-ACTCCTTCTGTGACTCCAGC-3’ and *IFN-γ* (5’-TTCTTCAGCAACAGCAAGGC-3’ and 5’-TCAGCAGCGACTCCTTTTCC-3’). The fold change for each transcript relative to β-actin housekeeping gene was calculated using the 2^-ΔΔCT^ method.

### Animals

Wild type C57BL/6, IL-17A–eGFP (*C57BL/6-Il-17a*^*tm1Bcgen*^*/J*) and IL-17A–KO (STOCK *Il-17a*^*tm1.1(ire)Stock*^*/J*) mice were obtained from Jackson Laboratory (Bar Harbor, ME). IL17A–eGFP mice contain an IRES-eGFP-SV40-polyA signal sequence cassette inserted after stop codon of *Il-17a* gene and express eGFP as a marker of IL-17A activity. Whereas IL-17A– KO mice contained abolished IL-17A expression due to insertional mutation of a codon optimized Cre-recombinase and a polyA signal into exon 1 of *Il-17a* gene. Groups of 4 to 6 mice at age 8-12 weeks old were used throughout the study. All mice were maintained in individually ventilated cages under specific pathogen free condition. Mice were euthanized with CO_2_ inhalation when they exhibited overt signs including hunched posture, fur ruffling, weakness, increased respiratory rate and difficulty breathing. This study has been approved by the Faculty of Medicine Ethics Committee for Animal Experimentation at the University of Malaya.

### *In vivo* infection

Fresh cultures of *C. neoformans* were washed and harvested by centrifugation at 1800 ×g for 10 min. Cells were adjusted to 10^7^ cells/ml in phosphate buffer saline (PBS) using a hemocytometer. Mice were first anesthetized with intraperitoneal injection of a mixture of ketamine (90 mg/kg) and xylazine (10 mg/kg) before inoculated with intranasal pipetting of 20 μl (2 × 10^5^ cells) yeast suspension. For survival study, each infected mouse was examined daily from 2 to 6 weeks post infection. For other study, mice were euthanized with CO_2_ inhalation at 28 days post-infection and serum was collected from blood. Lung was lavaged with 1.0 ml PBS and BALF was collected. Lung, mediatinal lymph nodes (mLN), spleen and brain were excised for further analysis.

### CFU count

Brain and lungs from the mice were excised, weighed and homogenized in 1 ml PBS using glass slides. A total volume of 20 μl of the serially diluted homogenates (at 10, 100 and 1000 folds) were plated on SDA plates in duplicates and cultured at 30°C for 48 hours. Fungal load was quantified using colony forming unit (CFU) per ml by calculating yeast colonies on each plate.

### Bioplex assay

Sera from each mouse were collected for measurement of cytokines using Bio-plex Pro Mouse Th17 assay (Bio-rad, CA, USA) which included the following cytokines: IL-17A, IL-17F, IL-21, IL-22, IL-23, IL-31, IL-33 and MIP-3α according to the manufacturer’s instructions. The Multiplex bead working solution was diluted from 25× stock solution beads and 50 μl of it was added into each well followed by 50 μl of sample. Each cytokine standards and samples were assayed in duplicate as provided by manufacturer. Samples with microbeads were incubated at room temperature on a magnetic microplate shaker for 30 minutes. After incubation, Bio-Plex detection antibody working solution was then added, washed 3× with Bio-Plex wash buffer and finally 1× streptavidin-PE was added before reading the plate on the Bio-Plex 200 system (Bio-Rad). Cytokine concentrations from each tissue homogenates were calculated based on each cytokine standard curve.

### Statistics

All statistical analyses were performed using GraphPad Prism 6. Analyses between groups were performed using Student’s *t*-test, whereby a *P* value of <0.05 was considered statistically significant.

## ACKNOWLEDGEMENTS

This work was supported by University of Malaya Research Grant (RG525-13HTM) and Fundamental Research Grant Scheme (FRGS, FP014-2017A). Movahed E. was supported by a postgraduate research fund (PPP) from University of Malaya.

## REFERENCES

1. May RC, Stone NR, Wiesner DL, Bicanic T, Nielsen K. 2016. Cryptococcus: from environmental saprophyte to global pathogen. Nat Rev Microbiol 14:106–17.

2. Mitchell TG, Perfect JR. 1995. Cryptococcosis in the era of AIDS--100 years after the discovery of Cryptococcus neoformans. Clin Microbiol Rev 8:515–48.

3. Jarvis JN, Harrison TS. 2007. HIV-associated cryptococcal meningitis. AIDS 21:2119–29.

4. Bojarczuk A, Miller KA, Hotham R, Lewis A, Ogryzko NV, Kamuyango AA, Frost H, Gibson RH, Stillman E, May RC, Renshaw SA, Johnston SA. 2016. Cryptococcus neoformans Intracellular Proliferation and Capsule Size Determines Early Macrophage Control of Infection. Sci Rep 6:21489.

5. Hayes JB, Sircy LM, Heusinkveld LE, Ding W, Leander RN, McClelland EE, Nelson DE. 2016. Modulation of Macrophage Inflammatory Nuclear Factor kappaB (NF-kappaB) Signaling by Intracellular Cryptococcus neoformans. J Biol Chem 291:15614–27.

6. Oliveira DL, Freire-de-Lima CG, Nosanchuk JD, Casadevall A, Rodrigues ML, Nimrichter L. 2010. Extracellular vesicles from Cryptococcus neoformans modulate macrophage functions. Infect Immun 78:1601–9.

7. Tan GM, Looi CY, Fernandez KC, Vadivelu J, Loke MF, Wong WF. 2015. Suppression of cell division-associated genes by Helicobacter pylori attenuates proliferation of RAW264.7 monocytic macrophage cells. Sci Rep 5:11046.

8. Ma H, Croudace JE, Lammas DA, May RC. 2006. Expulsion of live pathogenic yeast by macrophages. Curr Biol 16:2156–60.

9. Alvarez M, Casadevall A. 2006. Phagosome extrusion and host-cell survival after Cryptococcus neoformans phagocytosis by macrophages. Curr Biol 16:2161–5.

10. Johnston SA, May RC. 2010. The human fungal pathogen Cryptococcus neoformans escapes macrophages by a phagosome emptying mechanism that is inhibited by Arp2/3 complex-mediated actin polymerisation. PLoS Pathog 6:e1001041.

11. Nicola AM, Robertson EJ, Albuquerque P, Derengowski Lda S, Casadevall A. 2011. Nonlytic exocytosis of Cryptococcus neoformans from macrophages occurs in vivo and is influenced by phagosomal pH. MBio 2.

12. Davis MJ, Tsang TM, Qiu Y, Dayrit JK, Freij JB, Huffnagle GB, Olszewski MA. 2013. Macrophage M1/M2 polarization dynamically adapts to changes in cytokine microenvironments in Cryptococcus neoformans infection. MBio 4:e00264–13.

13. Arora S, Olszewski MA, Tsang TM, McDonald RA, Toews GB, Huffnagle GB. 2011. Effect of cytokine interplay on macrophage polarization during chronic pulmonary infection with Cryptococcus neoformans. Infect Immun 79:1915–26.

14. Eastman AJ, He X, Qiu Y, Davis MJ, Vedula P, Lyons DM, Park YD, Hardison SE, Malachowski AN, Osterholzer JJ, Wormley FL, Jr., Williamson PR, Olszewski MA. 2015. Cryptococcal heat shock protein 70 homolog Ssa1 contributes to pulmonary expansion of Cryptococcus neoformans during the afferent phase of the immune response by promoting macrophage M2 polarization. J Immunol 194:5999–6010.

15. Hill JO, Harmsen AG. 1991. Intrapulmonary growth and dissemination of an avirulent strain of Cryptococcus neoformans in mice depleted of CD4+ or CD8+ T cells. J Exp Med 173:755–8.

16. Huffnagle GB, Yates JL, Lipscomb MF. 1991. Immunity to a pulmonary Cryptococcus neoformans infection requires both CD4+ and CD8+ T cells. J Exp Med 173:793–800.

17. Mody CH, Lipscomb MF, Street NE, Toews GB. 1990. Depletion of CD4+ (L3T4+) lymphocytes in vivo impairs murine host defense to Cryptococcus neoformans. J Immunol 144:1472–7.

18. Snelgrove RJ, Edwards L, Williams AE, Rae AJ, Hussell T. 2006. In the absence of reactive oxygen species, T cells default to a Th1 phenotype and mediate protection against pulmonary Cryptococcus neoformans infection. J Immunol 177:5509–16.

19. Kawakami K, Koguchi Y, Qureshi MH, Kinjo Y, Yara S, Miyazato A, Kurimoto M, Takeda K, Akira S, Saito A. 2000. Reduced host resistance and Th1 response to Cryptococcus neoformans in interleukin-18 deficient mice. FEMS Microbiol Lett 186:121–6.

20. Decken K, Kohler G, Palmer-Lehmann K, Wunderlin A, Mattner F, Magram J, Gately MK, Alber G. 1998. Interleukin-12 is essential for a protective Th1 response in mice infected with Cryptococcus neoformans. Infect Immun 66:4994–5000.

21. Jain AV, Zhang Y, Fields WB, McNamara DA, Choe MY, Chen GH, Erb-Downward J, Osterholzer JJ, Toews GB, Huffnagle GB, Olszewski MA. 2009. Th2 but not Th1 immune bias results in altered lung functions in a murine model of pulmonary Cryptococcus neoformans infection. Infect Immun 77:5389–99.

22. Abe K, Kadota J, Ishimatsu Y, Iwashita T, Tomono K, Kawakami K, Kohno S. 2000. Th1-Th2 cytokine kinetics in the bronchoalveolar lavage fluid of mice infected with Cryptococcus neoformans of different virulences. Microbiol Immunol 44:849–55.

23. Voelz K, May RC. 2010. Cryptococcal interactions with the host immune system. Eukaryot Cell 9:835–46.

24. Muller U, Stenzel W, Kohler G, Werner C, Polte T, Hansen G, Schutze N, Straubinger RK, Blessing M, McKenzie AN, Brombacher F, Alber G. 2007. IL-13 induces disease-promoting type 2 cytokines, alternatively activated macrophages and allergic inflammation during pulmonary infection of mice with Cryptococcus neoformans. J Immunol 179:5367–77.

25. Piehler D, Stenzel W, Grahnert A, Held J, Richter L, Kohler G, Richter T, Eschke M, Alber G, Muller U. 2011. Eosinophils contribute to IL-4 production and shape the T-helper cytokine profile and inflammatory response in pulmonary cryptococcosis. Am J Pathol 179:733–44.

26. Szymczak WA, Sellers RS, Pirofski LA. 2012. IL-23 dampens the allergic response to Cryptococcus neoformans through IL-17-independent and -dependent mechanisms. Am J Pathol 180:1547–59.

27. Conti HR, Shen F, Nayyar N, Stocum E, Sun JN, Lindemann MJ, Ho AW, Hai JH, Yu JJ, Jung JW, Filler SG, Masso-Welch P, Edgerton M, Gaffen SL. 2009. Th17 cells and IL-17 receptor signaling are essential for mucosal host defense against oral candidiasis. J Exp Med 206:299–311.

28. Vautier S, Sousa Mda G, Brown GD. 2010. C-type lectins, fungi and Th17 responses. Cytokine Growth Factor Rev 21:405–12.

29. Chai LY, van de Veerdonk F, Marijnissen RJ, Cheng SC, Khoo AL, Hectors M, Lagrou K, Vonk AG, Maertens J, Joosten LA, Kullberg BJ, Netea MG. 2010. Anti-Aspergillus human host defence relies on type 1 T helper (Th1), rather than type 17 T helper (Th17), cellular immunity. Immunology 130:46–54.

30. Kleinschek MA, Muller U, Brodie SJ, Stenzel W, Kohler G, Blumenschein WM, Straubinger RK, McClanahan T, Kastelein RA, Alber G. 2006. IL-23 enhances the inflammatory cell response in Cryptococcus neoformans infection and induces a cytokine pattern distinct from IL-12. J Immunol 176:1098–106.

31. Angkasekwinai P, Sringkarin N, Supasorn O, Fungkrajai M, Wang YH, Chayakulkeeree M, Ngamskulrungroj P, Angkasekwinai N, Pattanapanyasat K. 2014. Cryptococcus gattii infection dampens Th1 and Th17 responses by attenuating dendritic cell function and pulmonary chemokine expression in the immunocompetent hosts. Infect Immun 82:3880–90.

32. Peck A, Mellins ED. 2010. Precarious balance: Th17 cells in host defense. Infect Immun 78:32–8.

33. Hardison SE, Wozniak KL, Kolls JK, Wormley FL, Jr. 2010. Interleukin-17 is not required for classical macrophage activation in a pulmonary mouse model of Cryptococcus neoformans infection. Infect Immun 78:5341–51.

34. Movahed E, Munusamy K, Tan GM, Looi CY, Tay ST, Wong WF. 2015. Genome-Wide Transcription Study of Cryptococcus neoformans H99 Clinical Strain versus Environmental Strains. PLoS One 10:e0137457.

35. Wozniak KL, Hardison SE, Kolls JK, Wormley FL. 2011. Role of IL-17A on resolution of pulmonary C. neoformans infection. PLoS One 6:e17204.

36. Chen S, Yan H, Zhang L, Kong W, Sun Y, Zhang W, Chen Y, Deng A. 2015. Cryptococcus Neoformans Infection and Immune Cell Regulation in Human Monocytes. Cell Physiol Biochem 37:537–47.

37. Shourian M, Ralph B, Angers I, Sheppard DC, Qureshi ST. 2017. Contribution of IL-1RI Signaling to Protection against Cryptococcus neoformans 52D in a Mouse Model of Infection. Front Immunol 8:1987.

38. Shoham S, Levitz SM. 2005. The immune response to fungal infections. British journal of haematology 129:569–582.

39. Snydman DR, Singh N, Dromer F, Perfect JR, Lortholary O. 2008. Cryptococcosis in solid organ transplant recipients: current state of the science. Clinical Infectious Diseases 47:1321–1327.

40. Wozniak KL, Ravi S, Macias S, Young ML, Olszewski MA, Steele C, Wormley Jr FL. 2009. Insights into the mechanisms of protective immunity against Cryptococcus neoformans infection using a mouse model of pulmonary cryptococcosis. PLoS One 4:e6854.

41. Jarvis JN, Casazza JP, Stone HH, Meintjes G, Lawn SD, Levitz SM, Harrison TS, Koup RA. 2013. The phenotype of the Cryptococcus-specific CD4+ memory T-cell response is associated with disease severity and outcome in HIV-associated cryptococcal meningitis. The Journal of infectious diseases 207:1817–1828.

42. Conti HR, Shen F, Nayyar N, Stocum E, Sun JN, Lindemann MJ, Ho AW, Hai JH, Jeffrey JY, Jung JW. 2009. Th17 cells and IL-17 receptor signaling are essential for mucosal host defense against oral candidiasis. Journal of Experimental Medicine 206:299–311.

43. Huang W, Na L, Fidel PL, Schwarzenberger P. 2004. Requirement of interleukin-17A for systemic anti-Candida albicans host defense in mice. Journal of Infectious Diseases 190:624–631.

44. Werner JL, Metz AE, Horn D, Schoeb TR, Hewitt MM, Schwiebert LM, Faro-Trindade I, Brown GD, Steele C. 2009. Requisite role for the dectin-1 β-glucan receptor in pulmonary defense against Aspergillus fumigatus. The Journal of Immunology 182:4938–4946.

45. Huang W, Na L, Fidel PL, Schwarzenberger P. 2004. Requirement of interleukin-17A for systemic anti-Candida albicans host defense in mice. J Infect Dis 190:624–31.

46. Bozza S, Zelante T, Moretti S, Bonifazi P, DeLuca A, D’Angelo C, Giovannini G, Garlanda C, Boon L, Bistoni F, Puccetti P, Mantovani A, Romani L. 2008. Lack of Toll IL-1R8 exacerbates Th17 cell responses in fungal infection. J Immunol 180:4022–31.

47. Zelante T, Bozza S, De Luca A, D’Angelo C, Bonifazi P, Moretti S, Giovannini G, Bistoni F, Romani L. 2009. Th17 cells in the setting of Aspergillus infection and pathology. Med Mycol 47 Suppl 1:S162–9.

48. Movahed E, Tan GM, Munusamy K, Yeow TC, Tay ST, Wong WF, Looi CY. 2016. Triclosan Demonstrates Synergic Effect with Amphotericin B and Fluconazole and Induces Apoptosis-Like Cell Death in Cryptococcus neoformans. Front Microbiol 7:360.

49. Tay ST, Chai HC, Na SL, Hamimah H, Rohani MY, Soo-Hoo TS. 2005. The isolation, characterization and antifungal susceptibilities of Cryptococcus neoformans from bird excreta in Klang Valley, Malaysia. Mycopathologia 159:509–13.

